# Data Driven Modelling of a Chemically Defined Growth Medium for *Cupriavidus necator* H16

**DOI:** 10.1101/548891

**Authors:** Christopher C. Azubuike, Martin G. Edwards, Angharad M. R. Gatehouse, Thomas P. Howard

## Abstract

*Cupriavidus necator* is a Gram-negative soil bacterium of major biotechnological interest. It is a producer of the bioplastic 3-polyhydroxybutyrate, has been exploited in bioremediation processes, and it’s lithoautotrophic capabilities suggest it may function as a microbial cell factory upgrading renewable resources to fuels and chemicals. It remains necessary however to develop appropriate experimental resources to permit controlled bioengineering and system optimisation of this microbial chassis. A key resource for physiological, biochemical and metabolic studies of any microorganism is a chemically defined growth medium. Here we use 1 mL micro-well cell cultures, automated liquid handling and a statistical engineering approach to develop a model that describes the effect of key media components and their interactions on *C. necator* culture cell density. The model is predictive and was experimentally validated against novel media compositions. Moreover, the model was further validated against larger culture volumes at 100 mL and 1 L volumes and found to correlate well. This approach provides valuable and quantifiable insights into the impact of media components on cell growth as well as providing predictions to guide culture scale-up.

## Introduction

The Gram-negative, non-spore forming soil bacterium, *Cupriavidus necator* H16, is of biotechnological importance principally due to its ability to accumulate >80% of its dry cell weight as polyhydroxyalkanoate (PHA) - a biodegradable polymer and an alternative to petroleum-based polymers (1, 2). The accumulation of PHA (specifically 3-polyhydroxybutyrate (3-PHB)) by the bacterium is a carbon conservation mechanism. When nitrogen, oxygen, or phosphorus becomes limiting to cell growth the bacterium diverts excess carbon to 3-PHB. This carbon store supports growth when conditions improve (3–6). *C. necator* H16 is also chemolithoautotrophic, with the ability to use CO_2_, and formate or H_2_ as carbon and energy sources to support cellular metabolism (7). In the absence of oxygen it can use alternative electron acceptors (NO_3_-and NO_2_-) to carry out anaerobic respiration by denitrification (8, 9). Metabolic engineering of *C. necator* H16 has been demonstrated through the introduction of pathways for the biosynthesis of alcohols (6, 7, 10, 11), fatty acids (12–14), alkanes (15) and enzymes (16) under both heterotrophic and autotrophic growth conditions. The production of branched chain alcohols by *C. necator* H16 through electricity powered cellular synthesis (i.e. electrosynthesis), demonstrates the value of this bacterium as a chassis capable of exploiting renewable feedstock for the biosynthesis of valuable products (7). More widely, the genus is known to encode genes facilitating metabolism of environmental pollutants such as aromatics and heavy metals making it a potential microbial remediator (8, 9, 17–19).

As with any bacterium, the development of *C. necator* as an industrial chassis requires appropriate tools for studying and engineering the organism. One of these resources is the availability of a characterised, chemically defined growth medium. Chemically defined media are important to enable experimental reproducibility, to reliably characterise the genetics of the organism, to determine genotype by environment interactions, and to facilitate fundamental research of bacterial physiology that underpins bioengineering efforts. While different chemically defined media have been deployed for the cultivation of *C. necator* H16 (3, 10–12, 20–22) there is no consensus regarding the components that are required, the concentration of each component, or how each component interacts to affect growth of the bacterium (SI Table 1).

Design of Experiments (DOE) is an iterative, empirical approach that systematically explores the relationship between input variables (factors) and output variables (responses). The approach yields a structured set of data that can be used to build statistical models employed in understanding or optimising system performance. These statistical models can be validated against prior knowledge, internal statitical methods or ultimately - by their ability to predict responses from new combinations of factors. Despite its early origins within the biological sciences, DOE is not a common-place method for life scientists. Increasing availability of laboratory automation and high throughput technologies may be changing this. DOE has found use in the optimisation of metabolic pathways (23, 24), cell-free protein synthesis reactions (25) and codon-use algorithms (26). It has been applied to the study of genotype-by-genotype and genotype-by-environment interactions in yeast (27) and in re-purposing enzyme activities (28).

Here we employ a statistical engineering approach to build a data driven model that can accurately predict *C. necator* H16 growth responses to a range of media formulations. The model highlighted different formulations that allow reproducible and robust growth of *C. necator* H16 with the minimal concentration of each component and allowed us to identify and understand interactions between components of the media. Finally, the model allowed the learning from the small scale (1 mL) to be applied at larger volumes (100 mL and 1 L). Understanding the impact of each component of a chemically defined medium on *C. necator* growth is a fundamental tool for controlled exploration of the biotechnological potential of this important bacterium.

## Results

### Identifying main ingredients in chemically-defined media that affect the growth of *C. necator*

A comparison of the literature for the use of defined media for *C. necator* growth identified variety in both the nature and concentrations of macroelements (C, N, P, Ca, Mg, and S) and trace elements (Cu, Zn, Fe and Mn) required for robust cell growth (Table SI 1). An initial scoping trial was carried out using fructose, glucose, glycerol or sucrose to identify a principle carbon source to be used for subsequent work, and to determine the range of concentrations to be tested. Four scoping trials were performed one each at low and high concentrations of media components, and two trials at the midpoint values between the two extremes. Cultures were grown in 1 mL volumes in a 48-well plate format for 120 h, at 30°C, 200 rpm. We confirmed that optical density (OD_600nm_) was an appropriate surrogate for determining *C. necator* cell growth (Fig. SI 1A). At the ranges tested glucose, glycerol and sucrose supported little or no growth of *C. necator* (Fig. SI 1B-D), whilst fructose supported high growth except at the lowest concentrations (Fig. 1). The OD_600nm_ for the two midpoint experiments demonstrated a peak at 72 h followed by a plateau. Growth obtained from optical density measurements at OD_600 nm_ correlated well with the number of viable cells (CFU/mL). From these scoping trails it was established that fructose would be our principle carbon source, OD_600nm_ was an appropriate measure of cell growth, growth assays in 1 mL volumes in a 48-well plate format were appropriate for subsequent experiments, and recording OD_600 nm_ at 72 h provides a good balance between measuring growth rate and peak culture density.

**Fig. 1.**
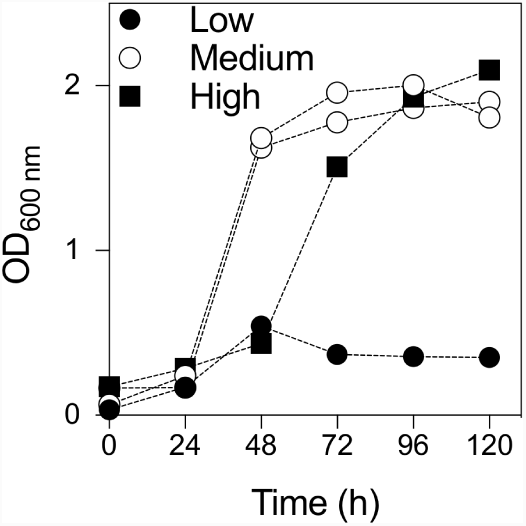
Scoping trials. Growth of *C. necator* H16 in 48-well format with fructose as the carbon source. Each experiment was carried out at low, medium (*n* = 2) or high concentrations of each media component (details can be found in Table S2).

We next identified key factors that might influence OD_600nm_ at 72 h. A definitive screening design (DSD) array was built based on 10 media components (Table 1). DSDs are highly efficient experimental designs in which all main effects can be estimated independently of other main effects and all possible two-way interactions. The requisite variant media compositions were assembled in a 48-well plate using a liquid-handling robot. Cell growth in these variant media was monitored as above. The resulting data were analysed using both the *Definitive Screening* and the *Two Level Screening* platforms in JMP Pro 13.0. Both analyses indicated that fructose, CaCl_2_ and amino acids contributed positively to growth, while Na_2_HPO_4_ and trace elements contribute negatively to growth. Factors such as NaH_2_PO_4_, K_2_SO_4_, MgSO_4_, NH_4_Cl and vitamins were not found to have statistically significant effects under the conditions tested (Fig. 2). These analyses also highlighted several two-way interactions that may influence growth responses, however, while DSDs are efficient arrays for de-aliasing main effects from other main effects and from second-order interactions, second-order interactions remain partially aliased with themselves. For this reason attributing effects to specific second-order interactions was not attempted at this stage.

**Fig. 2.**
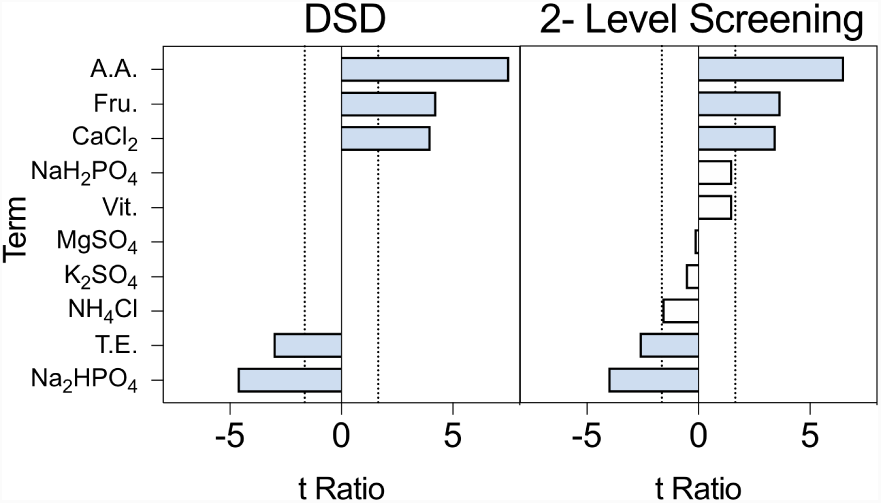
Definitive Screening Design array analysis. Definitive screening and 2-Level screening of data within DSD1 were performed. The comparative lengths of the t-ratios for each factor and factor interaction are shown. Bars extending to the right have a positive impact on the model, those extending to the left have a negative impact on the model. Terms deemed significant for model projection are highlighted (blue). Analysis is based on two replicated arrays (Table 1). The dashed vertical lines indicate a term that is significant at the 0.10 level.

**Table 1.** Definitive screening design array. A definitive screening design was developed to assess the impact of 10 ingredients found within the chemically defined media. All concentrations are in g/L except trace elements, amino acids and vitamins which are ml / L. Trace element working concentration contained (g/L): 15 FeSO_4_.7H_2_O, 2.4 MnSO_4_.H_2_O, 2.4 ZnSO_4_.7H_2_O, and 0.48 CuSO_4_.5H_2_O. A 100 *×* stock amino acid mix contained (g/L): 12.9 arginine, and 10 each of histidine, leucine and methionine. A 1000 *×* vitamin stock contained (g/L): 0.1 pyridoxine, 0.02 folic acid, 0.05 each of thiamine, riboflavin, niacin, pantothenic acid and nicotinamide. Abbreviations: Fru., fructose, T.E. Trace element mixture; A.A., amino acid mixture; Vit., vitamin mixture. The DSD was performed in replicate.

| Media | Fru. | NaH<sub>2</sub>PO<sub>4</sub> | Na<sub>2</sub>HPO<sub>4</sub> | K<sub>2</sub>SO<sub>4</sub> | MgSO<sub>4</sub> | CaCl<sub>2</sub> | NH<sub>4</sub>Cl | T.E. | A.A. | Vit. | OD1 | OD2 |
| --- | --- | --- | --- | --- | --- | --- | --- | --- | --- | --- | --- | --- |
| 1 | 20.25 | 0.1 | 0.1 | 0.01 | 0.01 | 0.01 | 0.01 | 0.01 | 0.05 | 0.01 | 1.06 | 1.37 |
| 2 | 20.25 | 6 | 6 | 4.8 | 2.8 | 0.8 | 4.8 | 4.8 | 19 | 4.8 | 1.14 | 1.62 |
| 3 | 0.5 | 3.05 | 0.1 | 4.8 | 0.01 | 0.01 | 0.01 | 4.8 | 19 | 4.8 | 0.52 | 0.47 |
| 4 | 40 | 3.05 | 6 | 0.01 | 2.8 | 0.8 | 4.8 | 0.01 | 0.05 | 0.01 | 0.09 | 0.78 |
| 5 | 0.5 | 6 | 3.05 | 0.01 | 2.8 | 0.01 | 0.01 | 0.01 | 19 | 4.8 | 0.77 | 0.71 |
| 6 | 40 | 0.1 | 3.05 | 4.8 | 0.01 | 0.8 | 4.8 | 4.8 | 0.05 | 0.01 | - | - |
| 7 | 40 | 6 | 0.1 | 2.405 | 2.8 | 0.01 | 4.8 | 4.8 | 19 | 0.01 | 0.93 | 0.69 |
| 8 | 0.5 | 0.1 | 6 | 2.405 | 0.01 | 0.8 | 0.01 | 0.01 | 0.05 | 4.8 | 0.11 | 0.28 |
| 9 | 0.5 | 6 | 0.1 | 4.8 | 1.405 | 0.01 | 4.8 | 0.01 | 0.05 | 0.01 | 0.80 | 0.64 |
| 10 | 40 | 0.1 | 6 | 0.01 | 1.405 | 0.8 | 0.01 | 4.8 | 19 | 4.8 | 1.62 | 1.48 |
| 11 | 0.5 | 6 | 6 | 0.01 | 2.8 | 0.405 | 0.01 | 4.8 | 0.05 | 0.01 | 0.29 | 0.32 |
| 12 | 40 | 0.1 | 0.1 | 4.8 | 0.01 | 0.405 | 4.8 | 0.01 | 19 | 4.8 | 1.86 | 1.41 |
| 13 | 0.5 | 6 | 6 | 4.8 | 0.01 | 0.8 | 2.405 | 0.01 | 19 | 0.01 | 0.41 | 0.55 |
| 14 | 40 | 0.1 | 0.1 | 0.01 | 2.8 | 0.01 | 2.405 | 4.8 | 0.05 | 4.8 | - | 0.01 |
| 15 | 0.5 | 0.1 | 6 | 4.8 | 2.8 | 0.01 | 4.8 | 2.405 | 0.05 | 4.8 | 0.17 | 0.43 |
| 16 | 40 | 6 | 0.1 | 0.01 | 0.01 | 0.8 | 0.01 | 2.405 | 19 | 0.01 | 1.69 | 1.67 |
| 17 | 0.5 | 0.1 | 0.1 | 4.8 | 2.8 | 0.8 | 0.01 | 4.8 | 9.525 | 0.01 | 0.17 | 0.28 |
| 18 | 40 | 6 | 6 | 0.01 | 0.01 | 0.01 | 4.8 | 0.01 | 9.525 | 4.8 | - | 0.19 |
| 19 | 0.5 | 0.1 | 0.1 | 0.01 | 2.8 | 0.8 | 4.8 | 0.01 | 19 | 2.405 | 0.48 | 0.70 |
| 20 | 40 | 6 | 6 | 4.8 | 0.01 | 0.01 | 0.01 | 4.8 | 0.05 | 2.405 | - | 0.04 |
| 21 | 20.25 | 3.05 | 3.05 | 2.405 | 1.405 | 0.405 | 2.405 | 2.405 | 9.525 | 2.405 | 1.76 | 1.70 |
| 22 | 0.5 | 0.1 | 6 | 0.01 | 0.01 | 0.01 | 4.8 | 4.8 | 19 | 0.01 | 0.86 | 0.71 |
| 23 | 0.5 | 6 | 0.1 | 0.01 | 0.01 | 0.8 | 4.8 | 4.8 | 0.05 | 4.8 | 0.10 | 0.19 |
| 24 | 40 | 6 | 0.1 | 4.8 | 2.8 | 0.8 | 0.01 | 0.01 | 0.05 | 4.8 | 1.64 | 1.56 |
| 25 | 40 | 0.1 | 6 | 4.8 | 2.8 | 0.01 | 0.01 | 0.01 | 19 | 0.01 | 0.20 | 0.23 |

### Augmentation of the data set

A definitive screening design can force many of the data-points collected to the edges of the design space. For this reason we ran a second DSD array (Table SI 3) in which the concentration ranges of the components were guided by data from DSD1 and the factors under investigation were restricted to those that were highlighted as significant in DSD1. Examining the combined data for DSD1 and DSD2 did indeed confirm that the extreme concentrations of some of the components were detrimental to cell growth. For example, when the concentrations of fructose were at the highest and lowest values (0.5 and 40 g/L) the OD_OD600nm_ 72 h were both lower and more variable than when fructose was restricted to between 5 and 25 g/L (Fig. 3). This indicates that maintaining fructose between 5 and 25 g/L is key to establishing robust and reliable growth. Likewise, adjustments were made to the concentration ranges of the amino acids (5 and 20 ml/L), CaCl_2_ (0.1 and 0.459 g/L) and Na_2_HPO_4_ (0.1 and 3.05 g/L). Other factors not identified as statistically significant were kept at the midpoint from DSD1, with the exception of NH_4_Cl which was set at the lowest value because there was some evidence of a negative impact from the 2-level factor analysis (Fig. 2).

**Fig. 3.**
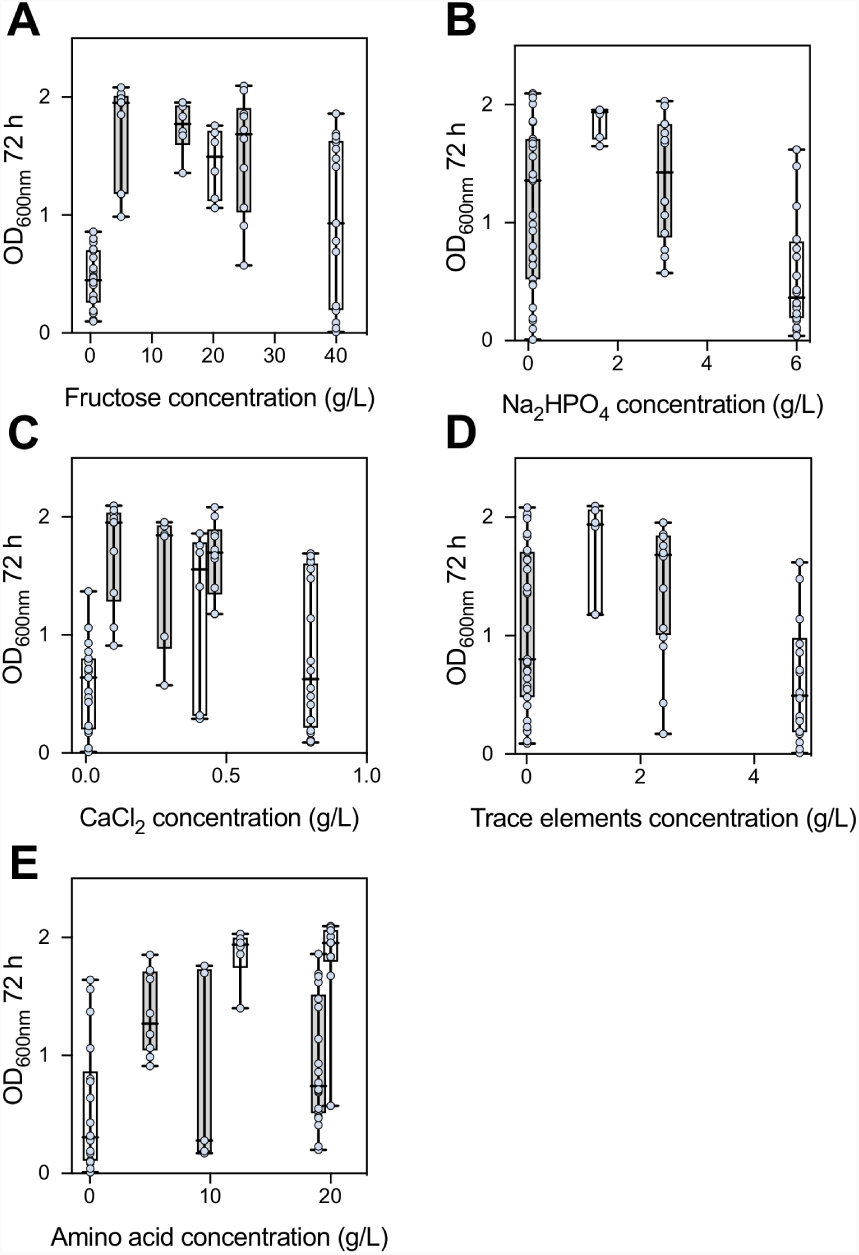
Combined data for DSD1 and DSD2 for key media components. **A.** Fructose. **B.** Na2HPO4 **C.** CaCl2 **D.** Trace elements **E.** Amino acids.

Examination of the distributions for DSD2 revealed that the media compositions that generated the highest cell densities (OD_600nm_ >2.0) were associated with low trace element concentrations and high amino acid content (Fig. SI 2). We therefore probed the putative interaction between amino acids and trace elements using a medium that was formulated to permit robust cell growth over 72 h. It was observed that the absence of trace elements did not adversely affect growth of *C. necator* but that the absence of amino acids did (Fig. 4A). Interestingly, simultaneous exclusion of both amino acids and trace elements resulted in cell densities comparable with the control. We hypothesized that in the absence of one or more of the amino acids (methionine, histidine, leucine and/or arginine), one or more of the components of the trace elements (CuSO_4_, FeSO_4_, MnSO_4_ and/or ZnSO_4_) inhibits growth of *C. necator*. This was tested first by formulating media without amino acids and withdrawing each trace element in turn (Fig 4B). Under these conditions, media without CuSO_4_ but containing other trace elements resulted in growth comparable to the control, while all three formulations that contained CuSO_4_ had reduced cell densities. These observations support the hypothesis that CuSO_4_ - in the absence of amino acids - inhibits *C. necator* growth. To determine which amino acid(s) interacts with CuSO_4_, a series of experiments were performed in which each amino acid was excluded in formulations with and without CuSO_4_. The first observation was that at high concentrations of amino acids the presence or absence of CuSO_4_ did not affect growth (Fig 4C). The second observation was that at lower concentrations of amino acids growth was partially suppressed in the presence of CuSO_4_ but that this was exacerbated in the absence of CuSO_4_. CuSO_4_ is therefore an important medium component and cannot simply be excluded from formulations. Similar growth responses were seen in experiments in which either methionine or leucine were excluded (Fig 4D and E). If arginine, or most notably histidine, were removed from the medium, then the presence of CuSO_4_ impaired growth of *C. necator* (Fig 4F and G). For media lacking histidine this effect was also observable when all other amino acid levels were kept high (Fig. 4G). From this data we conclude two points: first, that histidine protects against the inhibitory effects of copper, but second, CuSO_4_ is an essential component of the media whose absence retards growth when amino acid content is restricted. Both CuSO_4_ and histidine concentrations must therefore be balanced for robust growth.

**Fig. 4.**
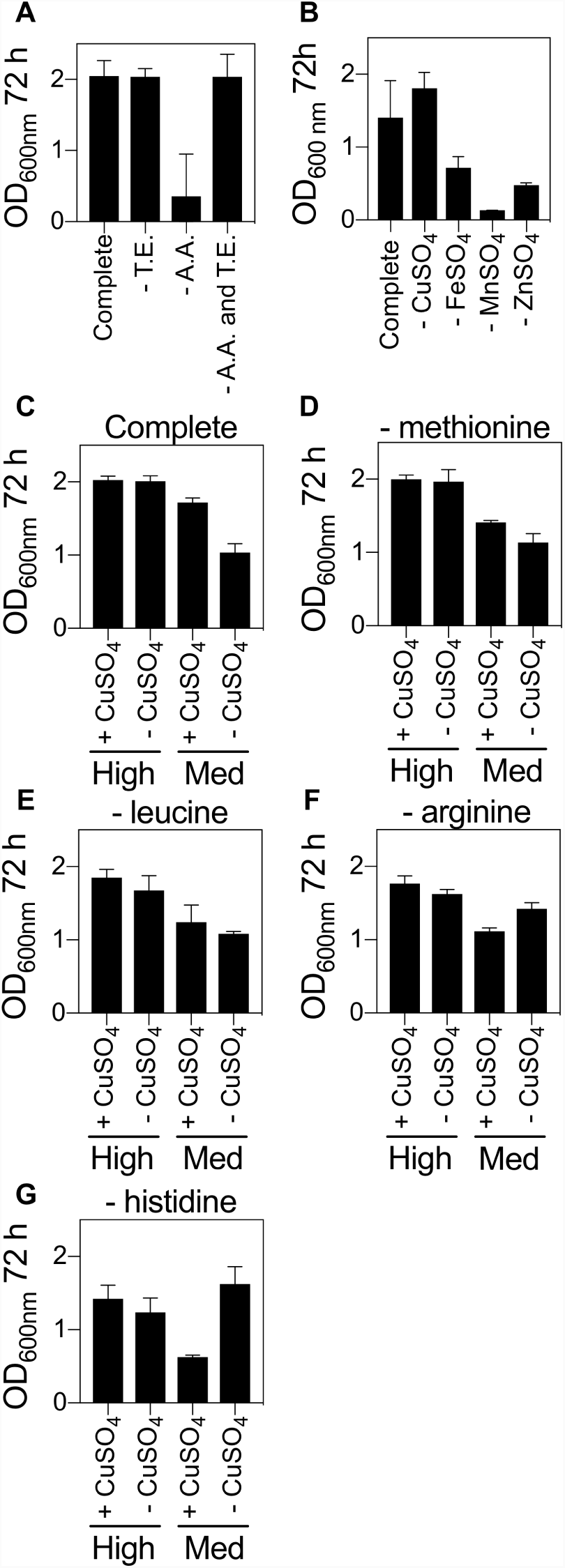
Interactions between trace elements and amino acids. **A.** *C. necator* grown in a complete, chemically defined medium with either amino acids, trace elements or both amino acids and trace elements excluded. **B.** *C. necator* grown in a complete medium with each of the four trace elements excluded. **C-G** *C. necator* grown in a complete, chemically defined medium with or without CuSO_4_ and with each of the four amino acids excluded. Error bars are S.E. Mean, *n* = 2.

### Final data augmentation

With a greater understanding of which components and concentrations are required to construct a medium supporting robust growth we re-investigated the roles that NaH_2_PO_4_, K_2_SO_4_, MgSO_4_ and NH_4_Cl have on the system. In the original DSD these factors were not identified as being statistically significant but those experiments were run under conditions in which key components (e.g. fructose and CaCl_2_ (Fig. 3) were at settings that have since been identified as resulting in poor or unreliable growth responses. We therefore wished to re-evaluated these factors under conditions in which Na_2_HPO_4_, CaCl_2_, trace elements and amino acid concentrations were not disruptive to cell growth (Table SI 4). Fructose was set at either 5 g/L or 25 g/L. The results indicated that these components did indeed affect growth rate when primary factors are not restricting growth (Fig. 5). Some of the suggestions from the original DSD were confirmed. Most notable was the observation that increasing NH_4_Cl had a detrimental affect on culture cell density. NaH_2_PO_4_ and K_2_SO_4_ had some detrimental impact if concentrations were low, whilst MgSO_4_ concentrations were not significant. These results were true at both high and low concentrations of fructose.

**Fig. 5.**
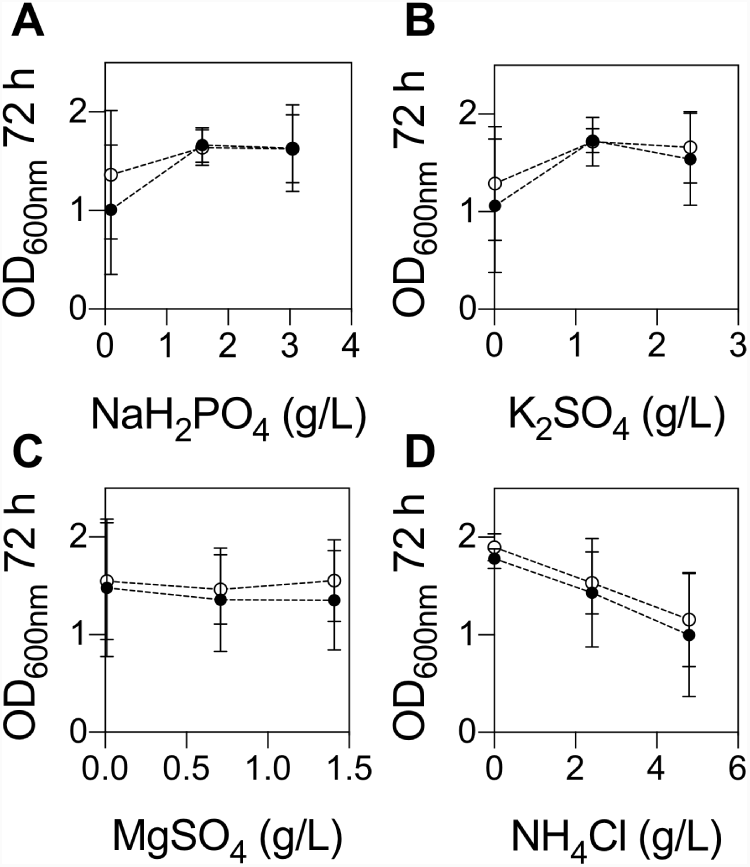
Re-examination of non-significant media components. The impact of components not deemed statistically significant in DSD1 were re-examined under less extreme conditions. These were **A.** NaH_2_PO_4_ **B.** K_2_SO_4_ **C.** MgSO_4_ and **D.** NH_4_Cl. Experiments were conducted at 5 g/L (open circles) or 25 g/L (closed circles) fructose. Error bars are S.E. Mean, *n ≥* 6

Finally, it is important to draw a distinction between what is statistically significant and what is essential. We observe little growth in formulae lacking MgSO_4_ yet MgSO_4_ is not deemed statistically significant. This is because under our experimental conditions, all concentrations of MgSO_4_ tested are in excess and do not limit to growth. There is therefore scope to reduce MgSO_4_ concentrations if this was desirable.

### Modelling the media formula-growth response landscape

At this stage, 64 different variant formulations have been experimentally assessed in duplicate across three different experiments (25, 13 and 26 experimental runs respectively). We then built a statistical model, trained against this data set, that could describe our understanding of how the cell cultures respond to changes in media composition and predict performance in novel formulations. We performed two-level screening on all 128 runs. This identified a number of factors and factor interactions deemed significant for model projection. The screening did not highlight fructose or K_2_SO_4_ (as these had been at held concentrations that did not significantly impact growth during much of DSDS2 and DSD3) but these terms were included manually in the model as we knew they were important factors. These terms were used to construct a standard least squares model. The least squares model was able to describe the relationships within the data with good accuracy (Fig. 6A). Though the model was internally consistent it was important to know if it could predict OD_600nm_ at 72 h in formulations it had not encountered during model training. We assessed 16 new formulations with sampling biased towards media formulations that were predicted to be in the top 25% of media performance. Each of these was assessed in triplicate and the resulting OD_600nm_ compared to predicted OD_600nm_ (Fig. 6B,C). As predicted by the model, all of the new formulations fell within the upper quartile of formula performance.

**Fig. 6.**
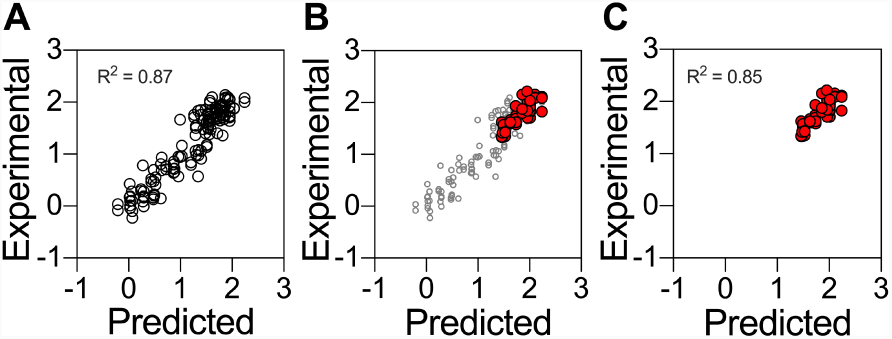
Experimental validation of model predictions in 48-well plate format. A. Model predicted values plotted against experimental data for least squares model. **B.** New experimental data for 1 mL culture volumes overlaid against the original least squares model. Open circles show the training data set, red circles are new data. **C.** New experimental data for 1 mL cultures plotted on their own against the model predicted values. Three replicates were assessed for each prediction.

The model allowed us to visualise phenomenon observed during the data-collection phase of the investigation. For example, it visualises the interaction between amino acids (specifically histidine) and trace elements (specifically copper) that was elucidated in Fig. 4. It shows that low and high concentrations of trace elements are detrimental to growth (low concentrations are to be avoided as these are essential to growth, while high concentrations may be toxic) and that increasing the amino acid concentration can help mitigate the inhibitory effects of high concentrations of trace elements (Fig. 7A). The greater the concentration of trace elements included the higher the amino acid concentration needs to be. Nevertheless, increasing the concentration of amino acids impacts in other areas. When fructose is low, increasing the concentration of amino acids increases cell density (Fig 7B). Given that the amino acids present in the media are the only alternative source of carbon for growth, raising the concentration of amino acids too high will impact on interpretation of experiments designed to examine carbon utilisation. A balance therefore needs to be struck between fructose, amino acids and trace element concentrations. The model also visualises interactions between Na_2_HPO_4_ and fructose (Fig. 7C). Greater concentrations of Na_2_HPO_4_ results in lower cell densities - an effect that can be partially offset by decreasing fructose concentrations. Finally, we had previously observed a negative effect of increasing NH_4_Cl concentrations (Fig. 5D). The model indicates that this can be mitigated by increasing the K_2_SO_4_ concentration (Fig. 7D). Understanding the interactions between media components is vital for predictions of culture performance and allows the experimenter to alter media formulations for different experimental goals.

**Fig. 7.**
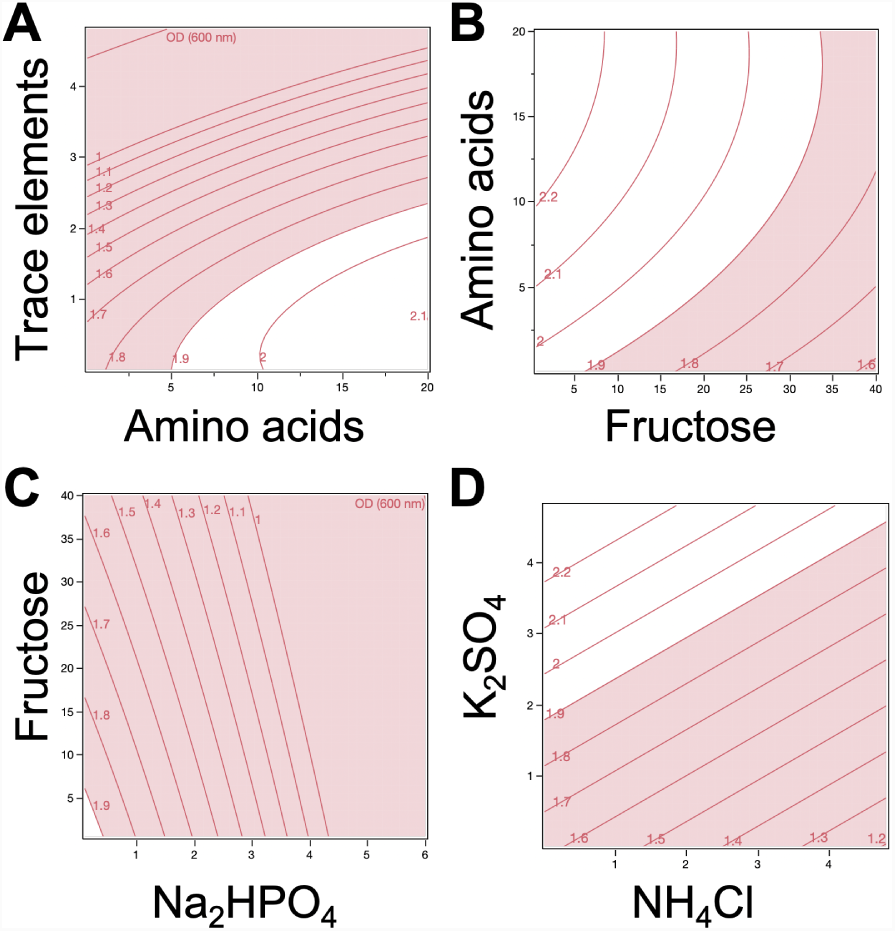
Interactions of components of the defined media described by the least squares model. **A.** Amino acid and trace element interactions. **B.** Fructose and amino acid interactions. **C.** Disodium phosphate and fructose interactions. **D.** Ammonium chloride and potassium sulphate interactions. Each panel represents a two way-interaction. Red zones are areas where OD_600nm_ at 72 h fails to reach 1.9, and white zones where it surpasses 1.9. Each contour grid line represents an OD_600nm_ of 0.1. All other media components were kept at concentrations permitting high growth.

### Model validation at greater volumes

We conducted two further model validation tests. We transferred these formulations to both 100 mL scale in shake flasks and 1 L batch fermentation in a bioreactor. Shake flask cultivations were carried out under identical conditions (30°C, 200 rpm) over 72 h periods in 250 mL baffled and non-baffled flasks, each containing 100 mL of medium. The formula for each cultivation was selected randomly from an L32 fractional factorial design of experiment (Fig SI 3). Growth (OD_600nm_) for each formula in both types of flasks were strikingly similar at every interval throughout the cultivation period. Most importantly, the growth rank for baffled and non-baffled flasks were identical. Spearman’s correlation showed a significant (p < 0.05) relationship between predicted and actual ranks for both flask types, with 0.867 correlation co-efficient *ρ* (Fig. 8A). Next, three formulae from shake flask cultivation together with two additional formulae were cultivated in 1 L bioreactor (Fig SI 4). Similarly, growth rank was predictable, with significant relationship between predicted rank and actual growth rank (p = 0.900) (Fig. 8B). During the culti-vations, there were no significant changes in bioprocess parameters. Constant agitation at 200 rpm, with 1 vvm airflow rate was sufficient to maintain dissolved oxygen (dO_2_) above 20%. Although no base was added in all cultivations, the pH of the media did not drop below 4.5, the set point. There was no significant relationship (*p* = 0.233; *p* = 0.520) between growth at 72 h and pH of media prior to inoculation or after growth.

**Fig. 8.**
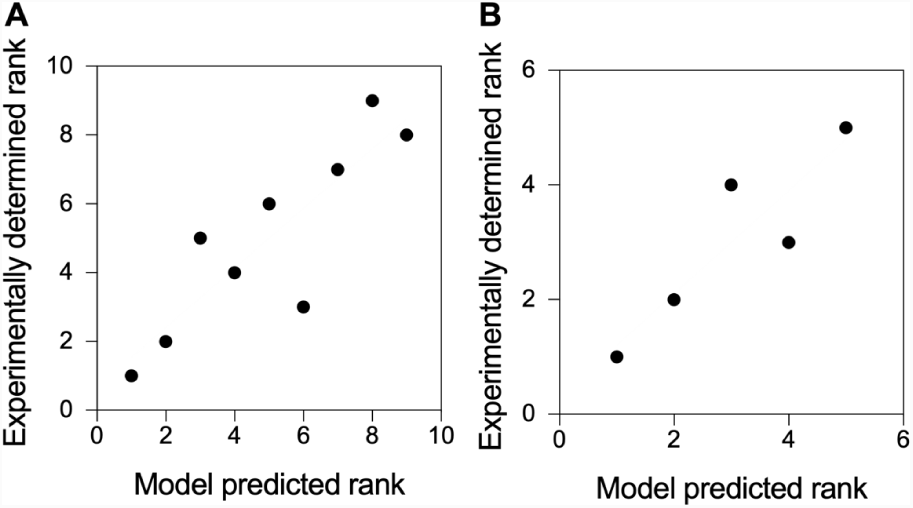
Experimental validation of model predictions at shake-flask and bioreactor scale. **A.** Spearman’s rank correlations between model predicted rank and experimental data for 100 mL culture volumes **B.** Spearman’s rank correlations between model predicted rank and experimental data for 1 L bioreactor batch flask cultures.

## Discussion

We developed a model trained against a structured data-set for the cultivation of *C. necator* H16 in a chemically defined medium with fructose as the primary carbon source. Our approach identified significant growth factors and their effects on culture density at 72 h. Fructose, glucose, glycerol and sucrose were used in the preliminary phase with fructose considered as the best carbon source supporting robust growth under heterotrophic conditions. While *C. necator* H16 has been reported to have broad substrate range its ability to utilize carbohydrates as a carbon source during heterotrophic growth appears to be limited to fructose and *N* acetylglucosamine (8, 17, 29). Utilisation of fructose by *C. necator* H16 is most likely to occur via substrate import via an ATP-binding cassette (ABC-type) transporter, followed by catabolism via 2-keto-3-deoxy-6-phosphogluconate (KDPG) and the Entner-Doudoroff pathway. The responsible genes, notably a putative regulator (*frcR*), ribosome transporters (i.e. *frcA, frcC* and *frcB* orthologs in *Escherichia coli* and *Ralstonia solanacearum*) and other essential genes facilitating such metabolism are located on chromosome 2 in-side gene clusters for glucose, 2-ketogluconate, and glucosamine catabolism (17). In contrast, phoshphofructokinase and 6-phosphogluconate dehydrogenase, key enzymes of the Embden-Meyerhoff-Parnas and oxidative pentose phosphate pathways, respectively, appear to be absent from the *C. necator* H16 genome. It is perhaps therefore not surprising that glucose supported little or no growth yet it has been reported that prolonged cultivation (> 70 h) with glucose as the sole carbon source resulted in a mutant that was able to utilize glucose (30). The mutant acquired such ability by mutating the *N*-acetylglucoasmine phosphotransferase system, which when deleted led to inability to utilize glucose (30, 31). Glycerol supported low growth of *C. necator* H16 especially in the medium forcing during the scoping trial. Such low growth was attributed to oxidative stress resulting from high levels of reactive oxygen species (ROS) formed as a result of elevated level of hydrogenases, leading to cell destruction by damaging DNA and other cell components (32).

Formulations of chemically defined media previously described for the cultivation of *C. necator* H16 are typically prepared without amino acids (3, 7, 11, 12, 22, 33). In this study, bacterial growth is improved when a small number of amino acids (arginine, histidine, leucine and methionine) are included in the medium. The data suggest that they serve as preferred sources of nitrogen, compared to NH_4_Cl. Their inclusion resulted in a shorter lag phase compared to media lacking amino acids. Although the effect of histidine is greater than methionine, arginine and leucine, their action was synergistic. Interestingly, the model indicated that there was an interaction between fructose and amino acid content that indicates the carbon : nitrogen ratio balance must be maintained, irrespective of the actual values. Growth reductions are less under low amino acid and low fructose concentrations (5 ml/L and 5 g/L, respectively) compared to that under low amino acid and high fructose concentrations (5 ml/L and > 20 g/L, respectively).

Metal ions, especially divalent cations of d-block transition metals, are important for bacterial growth where they act as metalloenzyme cofactors in living cells. Their presence in high concentrations however, tends to be detrimental to cells (34). Copper is known to play diverse structural and catalytic functions owing to its ability to exist either in a reduced (Cu+) state with affinity for thiol and thioether groups, or an oxidized (Cu2+) state with more likely coordination for oxygen or imidazole nitrogen group of amino acids including histidine (35). Bacteria have been reported to respond to high levels of copper by one of three families of metalloregulatory repressors: CopY, CsoR, and CueR. Under high concentration, an integral membrane Cu+ transporter (P1B-type ATPase) - characterised by histidine-rich domains, and conserved cytosine/histidine motifs within specific transmembrane domains - exports copper from the cytoplasm into the periplasm where in Gram-negative bacteria further detoxification and exportation are carried out by other related enzymes (35). Further, studies have reported that copper toxicity affected sugar (glucose) utilization in different microorganisms, and its growth inhibition on fungi (*Neurospora crassa* and *Saccharomyces cerevisiae*) was due to impaired/suppression of histidine biosynthesis (36, 37). In general, the toxic effect of copper appears to be enhanced under histidine limitation or impaired histidine biosynthesis, and is neutralized by addition of histidine.

This study provides insight into the impact of media formulations on growth and cell density of *C. necator* H16. Our growth model shows that the bacterium is able to grow to high cell density with minimal concentration of each growth component, provided the concentration of amino acids and trace elements are balanced. When cultivated under our formulae, growth peaked at 72 h and remained stationary after a further 20 h. Fructose, amino acids, and CaCl_2_, and interactions between amino acids and fructose, fructose and CaCl_2_, and amino acids and trace elements have significant positive effects on cell density, whilst Na_2_HPO_4_, and trace elements, and interactions between fructose and Na_2_HPO_4_, have negative effects. Addition of amino acids shortens the lag phase of growth and results in reproducible high cell density over repeated experiments. Besides fructose, only magnesium is considered essential for growth, whilst amino acids and K_2_SO_4_ are considered important. Thus, a defined medium supporting high cell density of *C. necator* H16 lacking some components (i.e. Na_2_HPO_4_, CaCl_2_, NH_4_Cl, and trace elements) can still result in high cell density comparable to formulae with all components. Both the information provided and the statistical approaches taken in this study will inform further efforts aimed at optimising other *C. necator* responses. This includes the biosynthesis of polyhydroxyalkanoate, platform chemicals, proteins and other products as well as future optimisation of growth under lithoautotrophic conditions with CO_2_ and molecular hydrogen serving as carbon and energy sources, respectively.

## Materials and Methods

### Bacterial strains

*Cupriavidus necator* H16 (DSM 428) was obtained from Deutsche Sammlung von Mikrooganismen und Zellkulturen GmbH, German Collection of Microorganisms and Cell Cultures (DSMZ). The bacterial strain was resuscitated on nutrient agar (peptone 5 g/L, and meat extract 3 g/L) according to the supplier’s instructions, and incubated at 30°C for 48 h.

### Chemicals

Carbon sources (glucose, fructose, glycerol and sucrose), salts (with exception of MgSO_4_.H_2_O and NH_4_Cl), trace metals, amino acids (histidine, leucine and arginine) and some vitamins (thiamine, niacin, pantothenic acid) were obtained from Sigma-Aldrich. The remainder of the vitamins, MgSO_4_.H_2_O, NH_4_Cl and methionine were obtained from Duchefa Biochemie B.V., BDH chemicals and Formedium, respectively.

### Preparation of media

Solutions of glucose, fructose, sucrose, vitamins, amino acid and each trace metal solution were filter sterilized through 0.22 *µ*m filter. Glycerol, NaH_2_PO_4_, Na_2_HPO_4_, MgSO_4_.H_2_O, NH_4_Cl, K_2_SO_4_, CaCl_2_.2H_2_O solutions were autoclaved. The trace solution was made of CuSO_4_.5H_2_O (dissolved in 0.1M HCl), FeSO_4_.7H_2_O (freshly prepared during each trace reconstitution from individual stock), MnSO_4_.H_2_O, and ZnSO_4_.7H_2_O. Amino acid stock solution contained: arginine, histidine, leucine, and methionine, while vitamin stock solutions contained: folic acid, niacin, nicotinamide, pantothenic acid, pyridoxine, riboflavin and thiamine. Subsequently, medium components were added from individual stock solutions except for trace elements which were added from reconstituted working solution. Forty-eight well plates were used in all trials, and medium reconstitution in each well was carried out using an automated liquid handling system (Eppendorf epMotion M5073). All stock solutions were prepared using water as a solvent, and were further diluted in sterile distilled water unless stated otherwise.

### Inoculate preparation and growth measurement

For each experiment, 48 h colonies from LB agar were washed twice in sterile distilled water and diluted to a working inoculate concentration in the range of 108 cfu/ml. The inoculate was further diluted 1:100 in wells containing medium. Plates were incubated at 30°C, 170 rpm. Optical density (OD) at 600 nm was measured every 24 h using a Varioskan LUXTM Multimode Microplate reader (Thermo Scientific).

### Batch cultivation in a bioreactor system

Large-scale cultivations were carried out in a batch mode with 1 L chemically defined media contained in 2 L capacity fermentors (Applikon ADI fermentation system). During the cultivations, pH was maintained above 4.5, temperature at 30°C, agitation at 200 rpm. Dissolved oxygen (dO_2_) was maintained above 20 % (1, 6) with airflow at 1 vvm (volume of air per volume of medium). No anti-foam agents or base were added during cultivation. Starter culture media were of the same formulations with that used in fermenters, and were prepared by growing 100 mL culture in 250 mL non-baffled flasks to late exponential growth at 30°C, 200 rpm. Following polarization and calibration of the dO_2_ probe, fermenters were inoculated with 10 mL (48 h) starter culture and cultivations were monitored on-line and off-line over 72 h. Samples were taken every 24 h for off-line OD_600nm_.

### Data analyses

Experimental designs were created using JMP Pro statistical software (version 13.0) and data from each experiment was analysed using the same software. Graphics were generated using GraphPad Prism 7.0.

## ACKNOWLEDGEMENTS

Christopher C. Azubuike is a Commonwealth Scholar (NGCA-2016-060) funded by Department for International Development (DFID), UK government. Manuscript prepared using a modified version the HenriquesLab template available via www.overleaf.com.

## Supporting Information

**Table SI 1.**
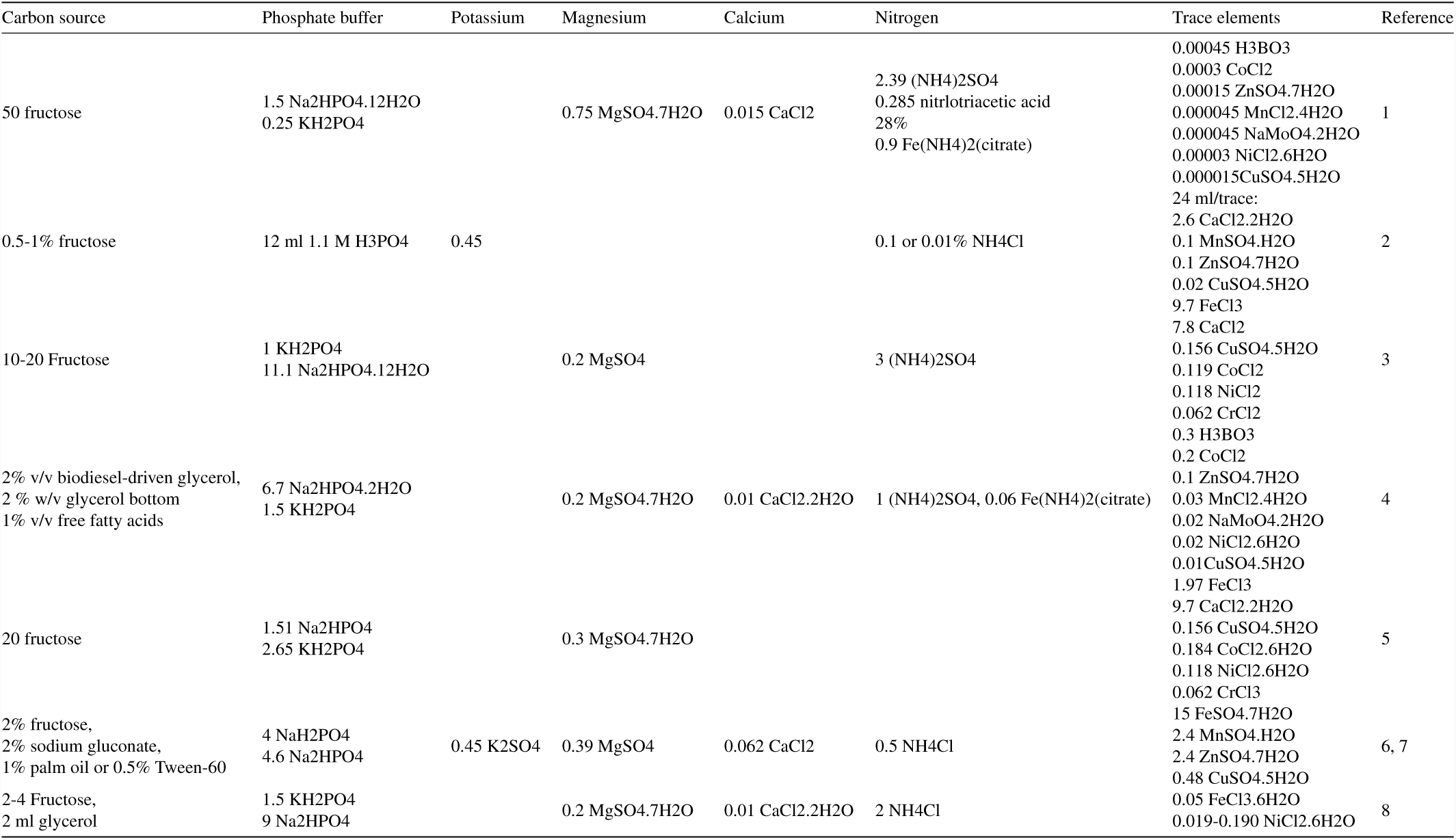
Previously described chemically defined media for the growth of *C. necator*. All concentrations are g/l unless otherwise indicated. References: 1. Grousseau et al. Metab Eng 42:74–84. 2. Müller et al. 2013. Appl Environ Microbiol. 3. Obruca et al. 1261–1267. 4. Sharma et al. AMB Express 6:36. 5. Aneja et al. Biotechnol Lett 31:1601–1612. 6. Lu et al. Appl Microbiol Biotechnol 96:283–297. 7. Lu et al. Appl Microbiol Biotechnol 97:2443–2454. 8. Oda et al 2013 Microb Cell Fact 12:2.

**Table SI 2.**
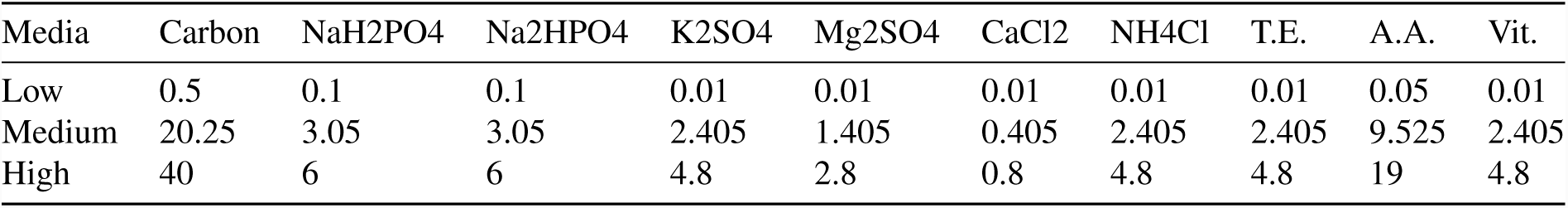
Scoping experiment. A scoping trial was developed to assess the impact of 10 basic ingredients found within the chemically defined media. All concentrations are in g/L except trace elements, amino acids and vitamins which are ml / L. Abbreviations: T.E. Trace element mixture; A.A., amino acid mixture; Vit., vitamin mixture. Carbon: fructose, glucose, glycerol or sucrose. The medium trial was performed in duplicate.

| Media | Carbon | NaH<sub>2</sub>PO<sub>4</sub> | Na<sub>2</sub>HPO<sub>4</sub> | K<sub>2</sub>SO<sub>4</sub> | Mg<sub>2</sub>SO<sub>4</sub> | CaCl<sub>2</sub> | NH<sub>4</sub>Cl | T.E. | A.A. | Vit. |
| --- | --- | --- | --- | --- | --- | --- | --- | --- | --- | --- |
| Low | 0.5 | 0.1 | 0.1 | 0.01 | 0.01 | 0.01 | 0.01 | 0.01 | 0.05 | 0.01 |
| Medium | 20.25 | 3.05 | 3.05 | 2.405 | 1.405 | 0.405 | 2.405 | 2.405 | 9.525 | 2.405 |
| High | 40 | 6 | 6 | 4.8 | 2.8 | 0.8 | 4.8 | 4.8 | 19 | 4.8 |

**Table SI 3.** Definitive Screening Design 2. A second definitive screening design was developed to assess the impact of the five ingredients identified as important to the chemically defined media. All concentrations are in g/L. Abbreviations with exception of T.E. and A.A, which are in ml/L: Fru., fructose, T.E., Trace element mixture; A.A., amino acid mixture. The DSD array was performed in duplicate.

| Media | Fru. | NaH<sub>2</sub>PO<sub>4</sub> | Na<sub>2</sub>HPO<sub>4</sub> | K<sub>2</sub>SO<sub>4</sub> | Mg<sub>2</sub>SO<sub>4</sub> | CaCl<sub>2</sub> | NH<sub>4</sub>Cl | T.E. | A.A. | OD1 | OD2 |
| --- | --- | --- | --- | --- | --- | --- | --- | --- | --- | --- | --- |
| 1 | 15 | 3.05 | 0.100 | 2.405 | 1.405 | 0.10 | 0.01 | 0.010 | 5 | 1.71 | 1.36 |
| 2 | 15 | 3.05 | 3.050 | 2.405 | 1.405 | 0.46 | 0.01 | 2.405 | 20 | 1.84 | 1.67 |
| 3 | 25 | 3.05 | 1.575 | 2.405 | 1.405 | 0.46 | 0.01 | 0.010 | 5 | 1.72 | 1.65 |
| 4 | 5 | 3.05 | 1.575 | 2.405 | 1.405 | 0.10 | 0.01 | 2.405 | 20 | 1.95 | 1.95 |
| 5 | 5 | 3.05 | 0.100 | 2.405 | 1.405 | 0.28 | 0.01 | 2.405 | 5 | 1.85 | 0.99 |
| 6 | 25 | 3.05 | 3.050 | 2.405 | 1.405 | 0.28 | 0.01 | 0.010 | 20 | 0.57 | 1.84 |
| 7 | 5 | 3.05 | 3.050 | 2.405 | 1.405 | 0.46 | 0.01 | 1.280 | 5 | 1.18 | 1.18 |
| 8 | 25 | 3.05 | 0.100 | 2.405 | 1.405 | 0.10 | 0.01 | 1.280 | 20 | 2.10 | 2.06 |
| 9 | 5 | 3.05 | 3.050 | 2.405 | 1.405 | 0.10 | 0.01 | 0.010 | 13 | 2.03 | 1.99 |
| 10 | 25 | 3.05 | 0.100 | 2.405 | 1.405 | 0.46 | 0.01 | 2.405 | 13 | 1.86 | 1.40 |
| 11 | 15 | 3.05 | 1.575 | 2.405 | 1.405 | 0.28 | 0.01 | 1.208 | 13 | 1.92 | 1.96 |
| 12 | 5 | 3.05 | 0.100 | 2.405 | 1.405 | 0.46 | 0.01 | 0.010 | 20 | 2.01 | 2.08 |
| 13 | 25 | 3.05 | 3.050 | 2.405 | 1.405 | 0.10 | 0.01 | 2.405 | 5 | 1.06 | 0.91 |

**Table SI 4.**
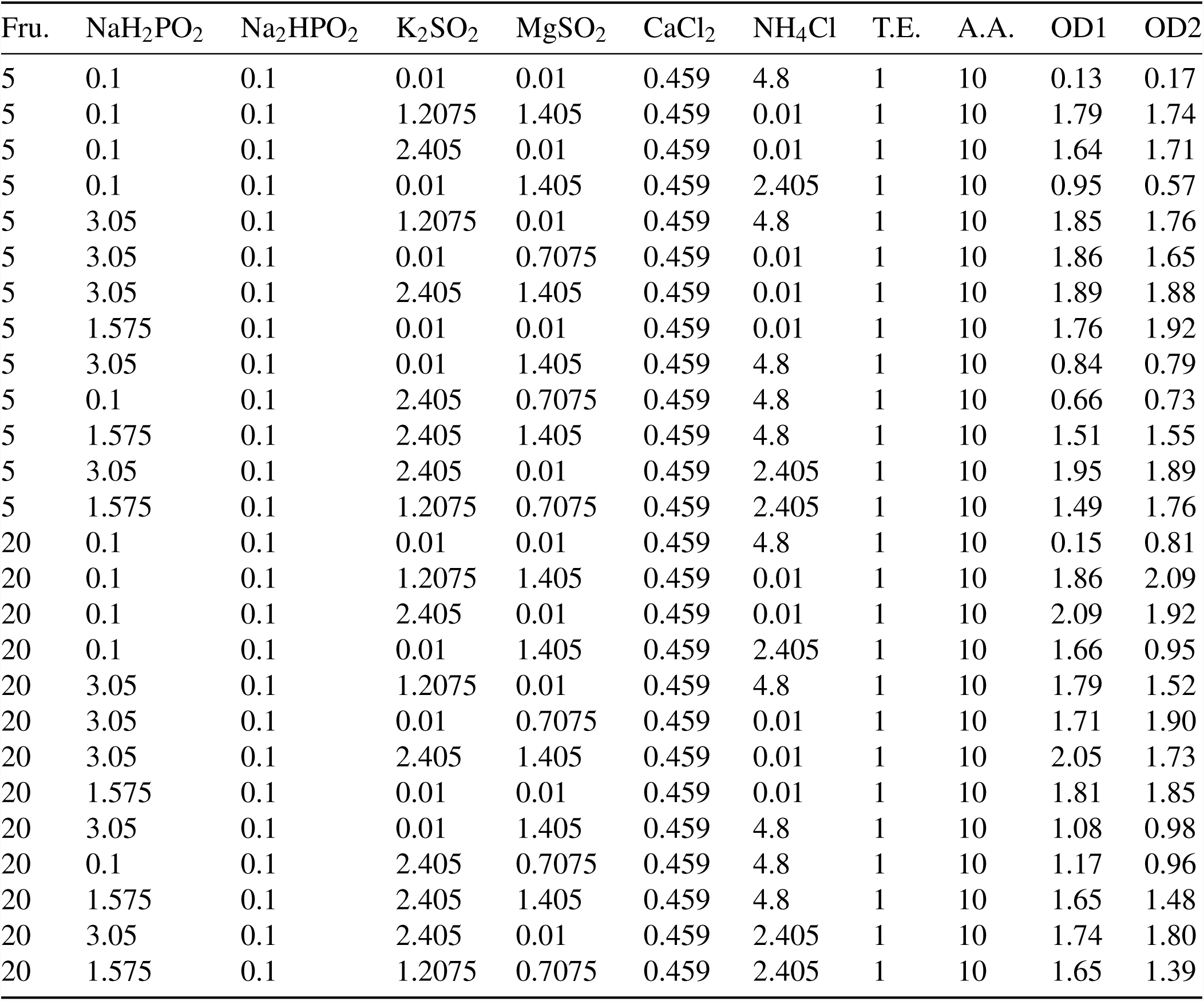
Definitive Screening Design 3. A final set of experiments were performed to re-assess the impact of components not deemed significant in the first DSD. All concentrations are in g/L. Abbreviations: Fru., fructose, T.E. Trace element mixture; A.A., amino acid mixture. The DSD array was performed in duplicate.

**Fig. SI 1.**
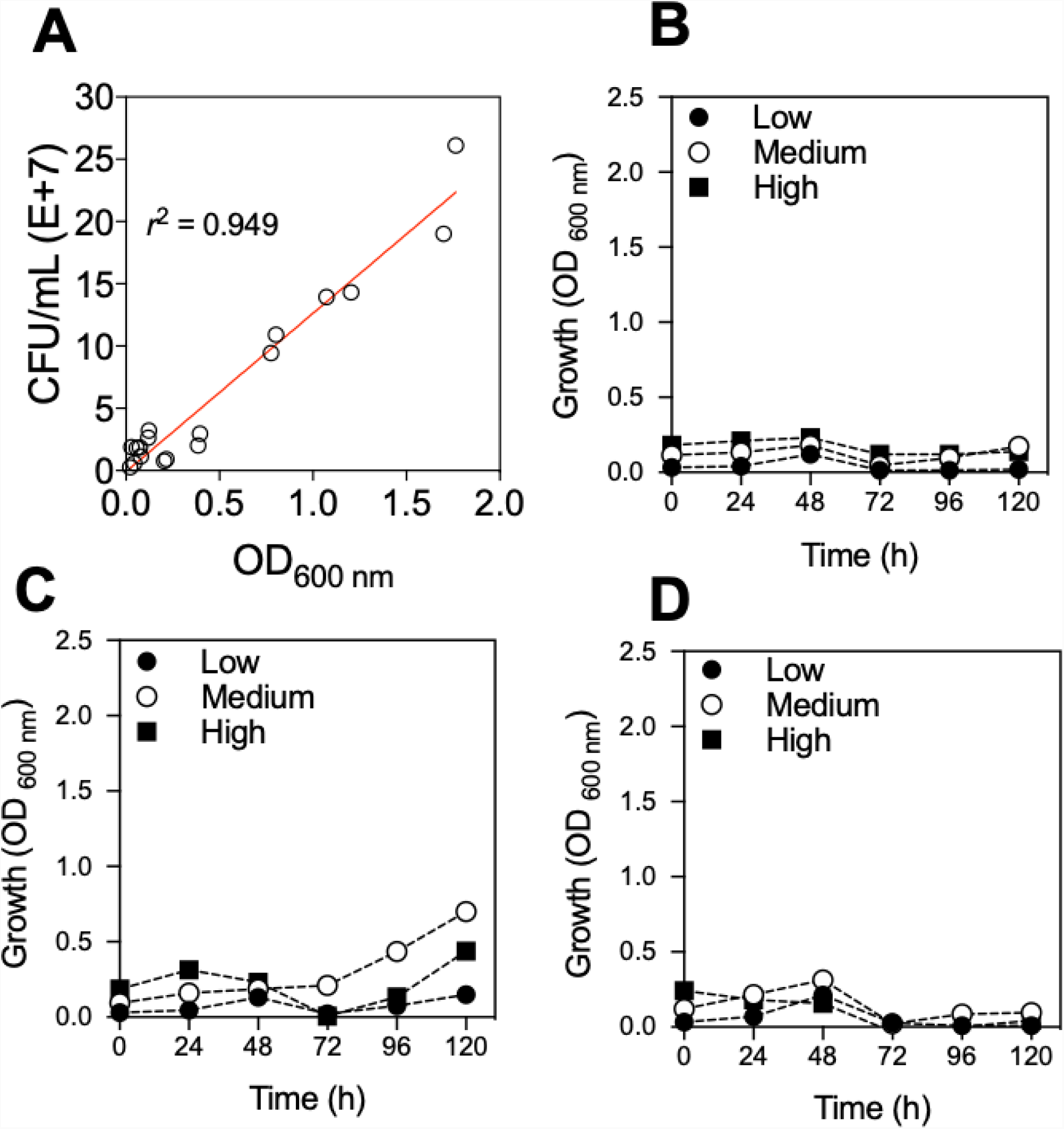
Growth characteristics of *C. necator* from initial scoping experiments. **A.** Colony forming units against OD_600nm_. Growth of *C. necator* on **B.** Glucose, **C.** Glycerol, and **D.** Sucrose as sole carbon source.

**Fig. SI 2.**
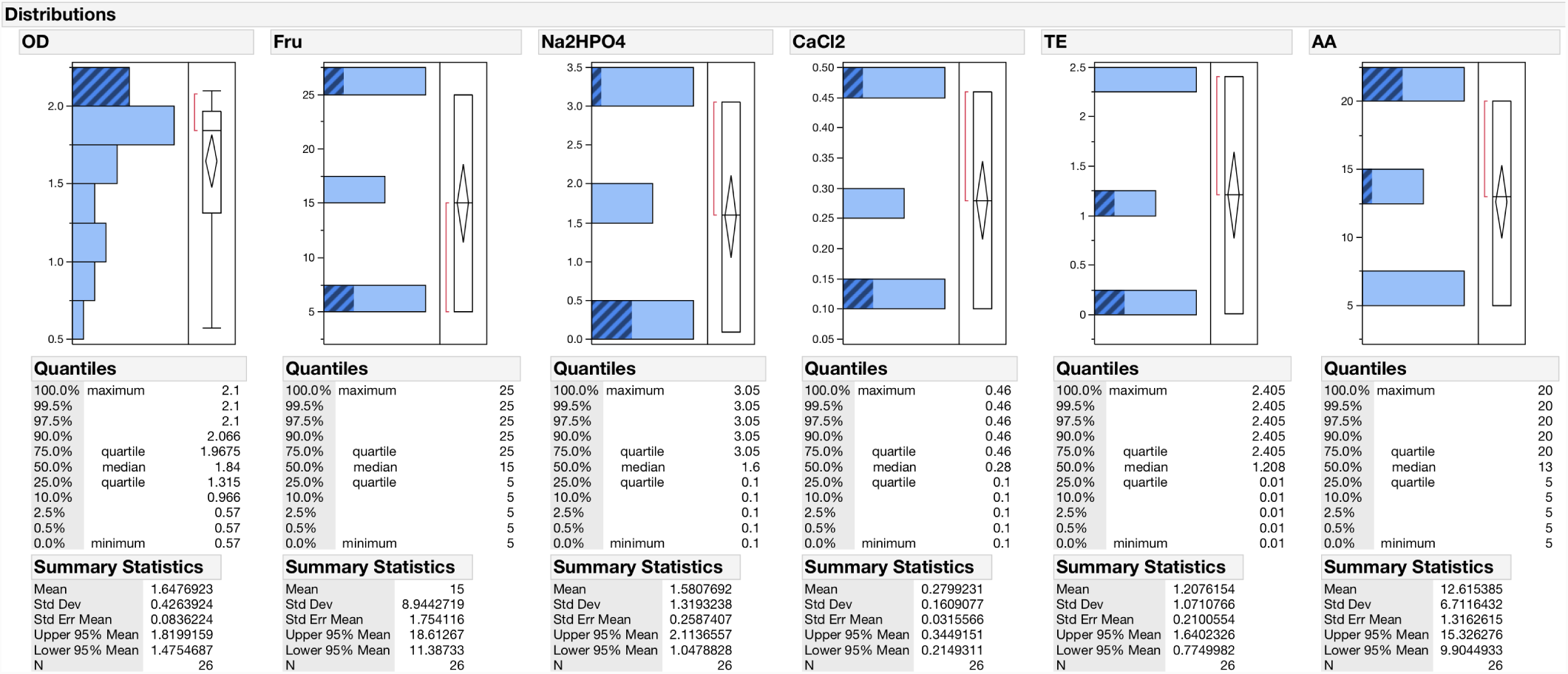
Distributions of data for Definitive Screening Design 2. Highlighted in blue cross-hatch are the settings that resulted in the greatest OD_600nm_ at 72 h for DSD2. Fructose, Na_2_HPO_4_ and CaCl_2_ may be at either the highest or lowest concentrations, whereas trace elements and amino acid concentrations are found at the lowest or highest concentrations respectively.

**Fig. SI 3.**
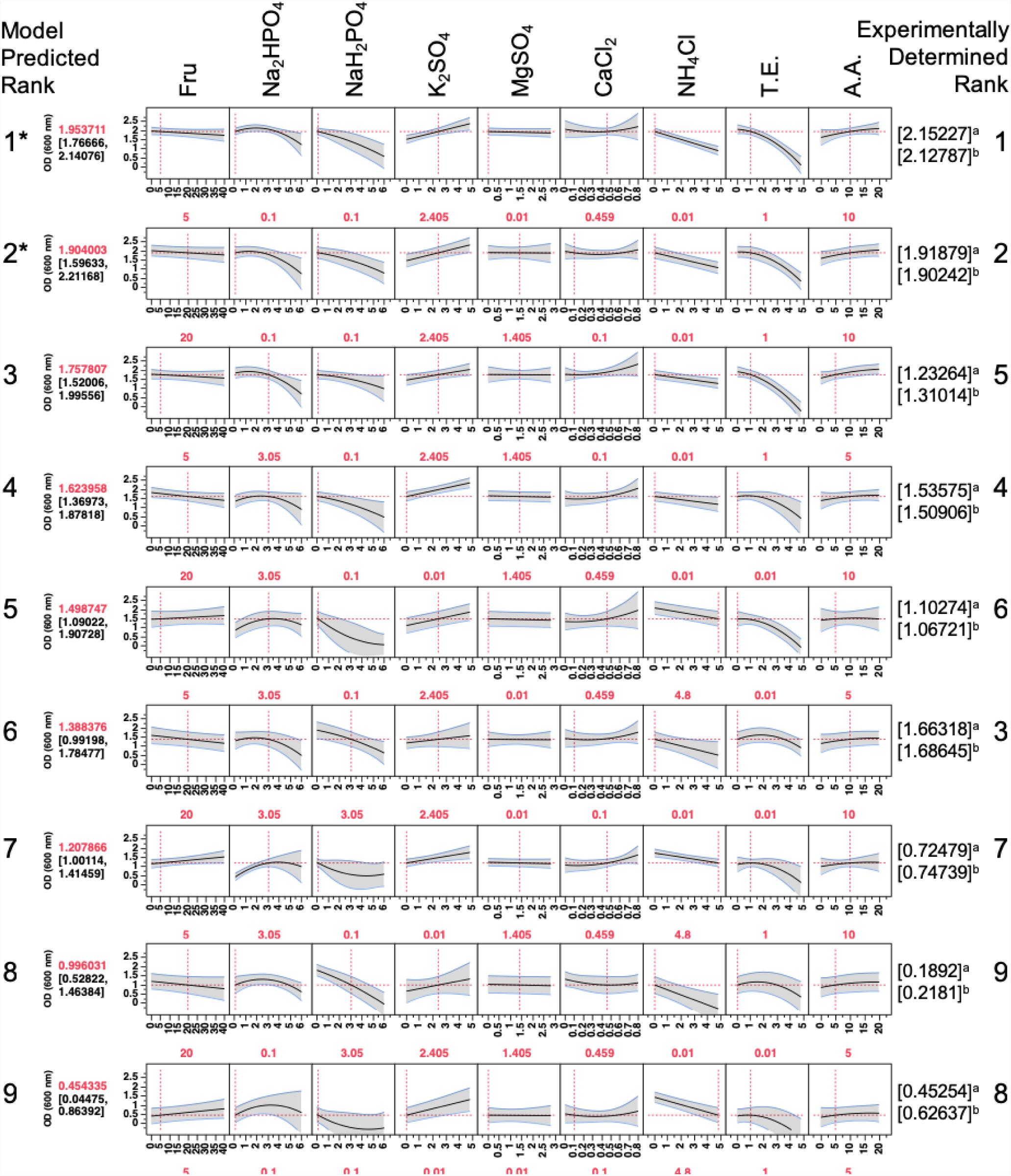
100mL validation. Nine different formulations were assessed at 100 mL culture volumes. For each row, the model predicted rank is shown on the left hand side, media settings for each component are shown in red underneath, the measured OD_(600nm)_ 72 h for two replicate experiments and experimentally determined rank on the right hand side.

**Fig. SI 4.**
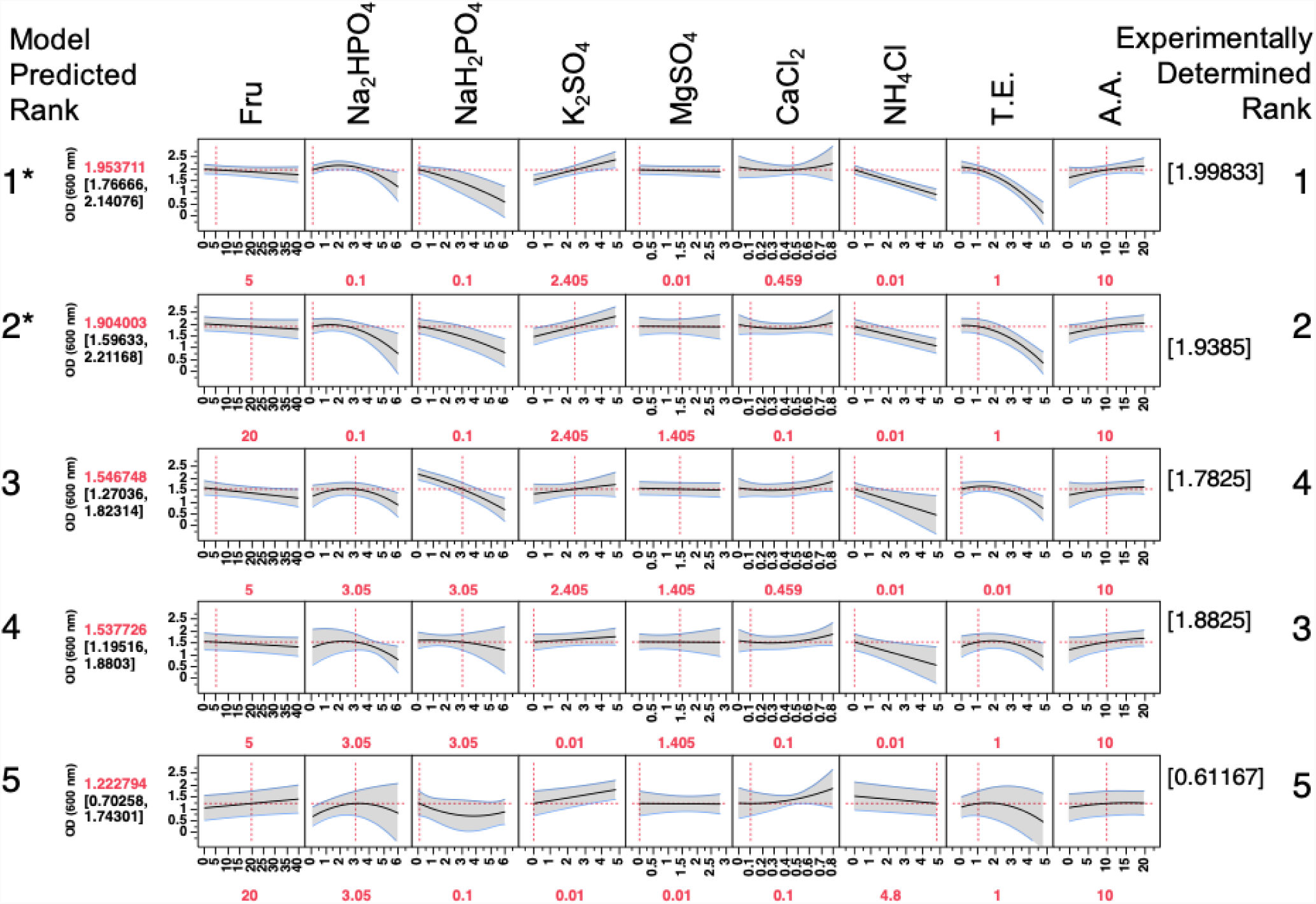
1 L validation. Five different formulations were assessed at 1 L culture volumes. For each row, the model predicted rank is shown on the left hand side, media settings for each component are shown in red underneath, the measured OD_(600nm)_ 72 h for two replicate experiments and experimentally determined rank on the right hand side.

